# Two loci contribute epistastically to heterospecific pollen rejection, a postmating isolating barrier between species

**DOI:** 10.1101/094912

**Authors:** Jennafer A. P. Hamlin, Natasha A. Sherman, Leonie C. Moyle

## Abstract

Recognition and rejection of heterospecific male gametes occurs in a broad range of taxa, although the complexity and redundancy of mechanisms underlying this postmating cryptic female choice is poorly understood. In plants, the arena for these interactions is the female reproductive tract (pistil), within which heterospecific pollen tube growth can be arrested via active molecular recognition. Unilateral incompatibility (UI) is one such pistil-mediated barrier in which pollen rejection occurs in only one direction of an interspecific cross. We investigated the genetic basis of pistil-side UI between *Solanum* species, with the specific goal of understanding the role and magnitude of epistasis between UI QTL. Using heterospecific introgression lines (ILs) between *Solanum pennellii* and *S. lycopersicum,* we assessed the individual and pairwise effects of three chromosomal regions (*ui1.1, ui3.1,* and *ui12.1*) previously associated with interspecific UI among *Solanum* species. Specifically, we pyramided *ui12.1* with each of *ui1.1* and *ui3.1*, and assessed the strength of UI pollen rejection in pyramided (double introgression) lines, compared to single introgression genotypes. We found that none of the three QTL individually showed UI rejection phenotypes, but lines combining *ui3.1* and *ui12.1* showed significant pistil-side pollen rejection. Furthermore, double introgression lines that combined different chromosomal regions overlapping *ui3.1* differed significantly in their rate of UI, consistent with at least two genetic factors on chromosome three contributing quantitatively to interspecific pollen rejection. Together, our data indicate that loci on both chromosomes 3 and 12 are jointly required for the expression of UI between *S. pennellii* and *S. lycopersicum* suggesting that coordinated molecular interactions among a relatively few loci underlying the expression of this postmating prezygotic barrier. In addition, in conjunction with previous data, at least one of these loci appears to also contribute to conspecific self-incompatibility, consistent with a partially shared genetic basis between inter- and intraspecific mechanisms of postmating prezygotic female choice.

## INTRODUCTION

Traits that underpin sexual recognition and rejection can be critical both for mate choice within species and for prezygotic isolating barriers between species. Such traits can contribute to premating interactions, including behavioral and chemical signals that indicate appropriate mating partners, or can act after mating but before fertilization, including interactions between gametes and/or between gametes and an internal reproductive tract. In the latter case, the female reproductive tract can be an important arena in which these interactions play out (Bernasconi *et al.*, 2004). Many species are known to exhibit “cryptic female choice” in which genotype-specific interactions between male gametes and female tissues determine the paternity of offspring following mating with more than one male genotype (Alonzo *et al.*, 2016). Similarly, female choice can influence the outcome of mating between species, when females are able to recognize and reject heterospecific male gametes. The specific mechanisms, by which this choice is exercised, either within or between species, have been identified in some select systems (Price, 1997; Manier *et al.*, 2013; Castillo and Moyle, 2014) and see also below). However, much remains to be understood about the complexity and redundancy of these postmating prezygotic female traits, the specific loci that are necessary and sufficient for recognition and rejection, and the extent to which these mechanisms are shared between intraspecific and interspecific sexual interactions.

Plants are among the many sexually reproducing organisms able to recognize and reject gametes from their own and other species. In angiosperms, recognition and rejection of pollen can occur at several stages after pollen is transferred (e.g. via wind or animal vectors) to the female receptive stigma (the receiving tissue for pollen deposition), including during pollen germination and pollen tube growth (via cell growth/elongation) down the female style (the reproductive tract that connect the stigma to the ovary). These ‘pollen-pistil’ interactions (the pistil is composed of the stigma and style) are roughly equivalent to post-copulatory interactions in animals, with the exception that pollen tubes (male gametophytes) actively express a substantial fraction of their own genome (Rutley and Twell, 2015). Molecular mechanisms of pistil-mediated recognition and rejection of conspecific pollen tubes are arguably best understood in the context of genetic self-incompatibility (SI), whereby pollen from self or close relatives is recognized and rejected in the female style. Intraspecific SI is mediated by the ‘S-locus’ which encodes (at least) two molecules responsible for the self-rejection mechanism: an *S-RNase* (the female/stylar component) that recognizes one or more pollen-expressed F-box protein(s) in germinated pollen tubes, and arrests pollen tube growth within the style. In gametophytic SI systems, pollen is rejected in styles when it bears an S-allele that is identical to an S-allele of the pistil (maternal) parent (McClure *et al.*, 1989). Because individuals will always share S-alleles with themselves, this genetic system prevents self-fertilization when pollen is transferred within a flower or between flowers on the same individual. In addition to genes at the S-locus, other factors are also known to be required for SI function, including HT—a small asparagine-rich protein (McClure *et al.*, 1999)—and other stylar glycoproteins (McClure *et al.*, 2000; Cruz-Garcia *et al.*, 2003; Hancock *et al.*, 2003; de Graaf, 1999) on the pistil-side, and pollen-side proteins including Cullins that are components of pollen protein complexes (Zhao *et al.*, 2010; Li and Chetelat, 2014; Hua and Kao, 2006). SI genotypes can give rise to SC lineages when one or more of these molecular components has a loss-of-function mutation(s) (Charlesworth and Charlesworth, 1979; Mable, 2008; Stone, 2002; Takayama and Isogai, 2005; Tao and Iezzoni, 2010; Covey *et al.*, 2010).

Pistil-mediated pollen tube rejection can also act as an important species barrier among plant lineages, especially those that are otherwise weakly isolated by trait differences associated with premating (e.g. pollinator or flowering time) isolation. In comparison to SI pollen-pistil interactions, the molecular basis of recognition and rejection of heterospecific pollen is less well understood. Nonetheless, in several cases there is strong evidence that elements of this behavior are mechanistically associated with SI. Classical studies of interspecific pollen-pistil barriers in some species groups indicate that these are often observed between lineages that differ in the presence/absence of SI, such that SI species styles reject pollen from self-compatible (SC) species, but the reciprocal cross does not show stylar rejection (the so-called “SI × SC rule”; de Nettancourt, 1997; Bedinger *et al.*, 2011; Lewis and Crowe, 1958; Murfett *et al.*, 1996). Molecular and functional analysis of the resulting “unilateral incompatibility” (UI) has confirmed that genetic components of SI can be shared in part with those of UI. For example, in some *Nicotiana* species, transforming non-rejecting SC genotypes with a functional *S-RNase* can be sufficient to confer the ability to reject pollen from other SC species (Hancock *et al.*, 2003). Additional factors required for SI can also play a role in stylar (female)-side UI. For example, both *S-RNase* and HT protein are required for SI *Nicotiana alata* rejection of pollen from SC *N. plumbaginifolia*; normal *S-RNase* expression is individually insufficient (Hancock *et al.*, 2003; McClure *et al.*, 1999). Similarly, in *Solanum* co-transformation of functional copies of both HT and *S-RNase* into SC species *S. lycopersicum* is sufficient to confer the ability to reject pollen from other SC species (Tovar-Méndez *et al.*, 2014).

While these observations clearly indicate that SI-associated molecular mechanisms can be sufficient to enable pistil-side pollen rejection between species, other data indicate that these mechanisms are more complex in nature. In particular, there is also evidence for *S-RNase-* independent UI mechanisms (Murfett *et al.*, 1996), often from species pairs that exhibit UI, but do not follow the SI × SC rule. For example, in *Solanum*, some SC populations of *S. pennellii* and *S. habrochaites* lack *S-RNase* expression but continue to be competent to reject interspecific pollen from SC species (Covey *et al.*, 2010; Baek *et al.*, 2015; Chalivendra *et al.*, 2013). Less directly, QTL mapping between *Solanum* species that both differ in SI-status and show UI, have detected UI QTL that do not colocalize with the S-locus. An analysis between SI *S. habrochaites* and SC *S. lycopersicum* detected three UI QTL, only one of which was localized to the S-locus (Bernacchi and Tanksley, 1997). A second study between SI *S. pennellii* and SC *S. lycopersicum* detected two UI QTL, neither of which was at the S-locus (Jewell, 2016). Interestingly, the two non-S-locus QTL detected in these studies (*ui3.1, ui12.1*) both localize to the same genomic regions on chromosomes 3 and 12, suggesting a common genetic basis for UI among different species. Moreover, in both studies, one of these QTL (*ui12.1*) co-localized with the known genomic location of HT, and in one case (Jewell, 2016) the presence/absence of HT expression in mature styles was significantly associated with the phenotypic strength of UI.

Overall, these data suggest that there might be substantial overlap between molecular mechanisms of SI and UI, including both *S-RNase*-dependent or -independent mechanisms, but also involvement of additional loci that have yet to be molecularly identified, and whose relationship to SI is unknown. As such, several important aspects of the genetics of pistil-side UI remain unclear, including the minimum number of factors sufficient to express *S-RNase*-independent UI, the degree of overlap with molecular loci underpinning *S-RNase*-dependent mechanisms, and therefore the level of redundancy between alternative mechanisms underlying these important postmating forms of female mate choice.

Here our goal was to assess the specific role of three chromosomal regions in affecting pistil-side UI between species. Focusing on the UI QTL previously identified in two different Solanum species crosses—*ui1.1, ui3.1*, and *ui12.1* (Bernacchi and Tanksley, 1997; Jewell, 2016)—our aim was to evaluate the individual and joint effects of these three unlinked chromosomal regions on the expression of UI. To do so, we used near isogenic lines (NILs), in which single chromosomal regions from a donor species genotype (*Solanum pennellii*) are introgressed into the genetic background of an otherwise isogenic recipient species (*Solanum lycopersicum*). Lines incorporating three different introgressed regions (from chromosomes 1, 3, and 12) were examined individually and in pairwise combinations; the latter double introgression lines (DILs: Canady *et al.* 2006; sometimes called ‘pyramid lines’: Gur and Zamir, 2004) were created via crosses among NILs (see Methods). Two criteria were used to evaluate evidence for epistasis between these target loci. First, we examined evidence for transmission ratio distortion (TRD) in the products of crosses between different NILs, to look for evidence that particular genotypes were over- or underrepresented. Second, we quantified the strength of pistil-side UI response phenotypes in the DIL lines and compared this to the same phenotype in single (NIL) introgression genotypes. This comparison allowed us to evaluate whether the quantitative effects of individual introgressions differs in the presence of a second introgressed locus, and to evaluate the minimum number of loci required to express pistil-side interspecific pollen rejection. We find evidence that loci on chromosomes 3 and 12 are simultaneously required in order to express UI; lines in which these loci are represented individually are unable to reject heterospecific pollen. One of these QTL—*ui12.1*—likely involves a known molecular contributor to SI (HT protein) thus further supporting the inference that factors associated with SI contribute to the expression of the UI phenotype. In addition, comparisons among DILs created using overlapping sections of *ui3.1* suggest that this other QTL might be underpinned by at least two separate genetic factors that additively contribute to the genetic variation in the strength of UI.

## METHODS

### Study System

The tomato clade (Solanum section Lycopersicon) contains 13 closely-related species known to be separated by a range of incomplete pre- and postzygotic isolating barriers (Moyle, 2008), including pollen-pistil incompatibility, and specifically UI (Bedinger *et al.*, 2011; Covey *et al.*, 2010). Baek *et al.* (2015) directly examined the strength of pollen tube rejection between all 13 species within the tomato clade and found evidence that UI was strongest and most consistently observed between SI × SC species. Among other genetic mapping resources in this group are several introgression line libraries in which chromosomal regions representing most or all of a donor species genome have been serially introgressed into the genetic background of a recipient species, usually the domesticated tomato *S. lycopersicum* (Bernacchi and Tanksley, 1997; Eshed and Zamir, 1995). For this study, we used lines drawn from a NIL library previously developed between *S. pennellii*, a wild tomato species, and *S. lycopersicum*, where each line contains a marker delimited homozygous region of *S. pennellii* accession LA0716 introgressed into the genomic background of *S. lycopersicum* accession LA3475 (Eshed *et al.*, 1992; Eshed and Zamir, 1995; Eshed and Zamir, 1994). We used four different lines drawn from this library (Supplemental Table 1). IL1-1 overlaps the genomic location of the *S*-locus as well as the location of *ui1.1*, the UI locus previously mapped in an F2 population between *S. lycopersicum* and *S. habrochaites* (Bernacchi and Tanksley, 1997). IL3-3 and IL3-4 contain *S. pennellii* introgression regions that overlap the previously mapped *ui3.1* in (Bernacchi and Tanksley, 1997) and in (Jewell, 2016), which together spans a broad genomic region (~70 cM); these lines contain an overlapping region of approximately 44 cM (Supplemental Figure 1). IL12-3 overlaps the previously mapped *ui12.1* in prior studies (Bernacchi and Tanksley, 1997).

Note that the known chromosomal location of HT protein falls within both IL12-3 and *ui12.1.* HT was duplicated in the ancestor of Solanum, resulting in two tandemly arrayed paralogs (HT-A and HT-B) at this chromosome 12 location (Covey *et al.*, 2010). Moreover, both HT-A and HT-B are expressed in *S. pennellii* LA0716 but are non-functional in *S. lycopersicum* due to null mutations in both HT-A and HT-B (Kondo *et al.*, 2002; Covey *et al.*, 2010). In contrast, the *S-RNase* protein at the S-locus is non-functional in both *S. pennellii* and *S. lycopersicum* genotypes in this experiment; *S. pennellii* is normally an SI species, but SI has recently been lost in this population due to a complete loss of the *S-RNase* gene (Solyc01g055200) (Li and Chetelat, 2015). Therefore we included an IL spanning *ui1.1* here in order to assess whether other genes contained within this chromosomal region also contribute to S-RNase-independent mechanisms of UI.

### Construction of Double Introgression Lines

Seeds for our four target introgression lines were obtained from the Tomato Genetics Resource Center (tgrc.ucdavis.edu). To generate lines with two introgressed *S. pennellii* regions (Double Introgression Lines, or DILs), crosses were performed pairwise between NIL lines (Table 1), and resulting heterozygous F_1_s were then selfed to generate F_2_ seeds (a ‘DIL population’) for genotyping and ultimately phenotyping. In this experiment, three different DIL combinations were generated (Table 1) in which we combined *ui12.1* with *ui1.1* and with the two alternative (overlapping) *S. pennellii* regions at *ui3.1.* We were unable to generate offspring from reciprocal crosses in all pairwise NIL-NIL combinations, as some of these did not produce seeds in one of the crossing directions despite numerous attempts, or failed to produce viable seed that was homozygous for target *S. pennellii* introgressions across each target QTL region. Patterns of marker representation and segregation distortion in F2 progeny from the crosses are discussed further below.

**Table 1.** Proportion of pollen tube growth for each genotype with upper (upr) and lower (lwr) 95% confidence intervals.

| **Genotype**<br>**(Female x Male)** | **Mean ± SE** | **CI.95.lwr** | **CI.95.upr** | **# of biological**<br>**replicates** |
| --- | --- | --- | --- | --- |
| **NIL 1-1** | 0.9398 ± 0.0154 | 0.8650 | 1.0144 | 3 |
| **NIL 3-3** | 0.9369 ± 0.0328 | 0.8621 | 1.011 | 3 |
| **NIL 3-4** | 0.9233 ± 0.0402 | 0.8681 | 1.002 | 4 |
| **NIL 12-3** | 0.9785 ± 0.0117 | 0.9206 | 1.036 | 5 |
| **DIL 1-1 x 12-3** | 0.8459 ± 0.0200 | 0.7811 | 0.9105 | 4 |
| **DIL 12-3 x 3-3** | 0.5082 ± 0.0317 | 0.4693 | 0.5987 | 4 |
| **DIL 3-4 x 12-3** | 0.7609 ± 0.0472 | 0.7080 | 0.8136 | 6 |

### Genotyping and Scoring Individuals

Progeny were genotyped within each F2 ‘DIL population’ in order to identify individuals that were homozygous for each *S. pennellii* introgression region and to describe patterns of marker transmission ratio distortion (TRD) at these introgressions. These genotypes were used to identify individuals that were homozygous at both introgression regions, for further phenotypic assessment. Genotypes were also used to calculate overall genotype frequencies in segregating populations, to assess if there was evidence of non-Mendelian patterns of transmission that might be consistent with selection against certain genotypic combinations.

Genomic DNA was extracted using a modified CTAB protocol and Cleaved Amplified Polymorphic Sequence (CAPS) genotyping was used to characterize the allelic identity (S. *lycopersicum:* L or *S. pennellii:* P) using the target markers (Supplemental Table 2). CAPS markers are restriction fragment variants caused by single nucleotide polymorphisms or insertion/deletions, which create or abolish restriction enzyme recognition sites. Here we identified and genotyped markers that were designed to distinguish *S. lycopersicum* and *S. pennellii* alleles.

For each individual, DNA was amplified using PCR primers for each target marker and checked with gel electrophoresis. For each individual at each marker locus, a subsample of the amplicon was incubated with the relevant restriction endonucleases, and the digestion products were separated on 1.5% agarose gels, visualized with Ethidium Bromide staining, and imaged prior to manual scoring. Markers were chosen such that an allele from one parent would yield unique sized fragments (in bp) and the allele from the other parent would yield fragments of a different size (Supplemental Table 2). Thus each F_2_ individual could be scored as homozygous for a parental allele (either *S. lycopersicum* or *S. pennellii*) at each marker, or as heterozygous in the case where the sample had the cleaved banding patterns representative of both parental alleles (Supplemental Table 2).

Each F_2_ individual was genotyped at 2-4 markers associated with the chromosomal region they were expected to carry. For each target introgression on chromosomes 1 and 12, three markers were selected to span the length of the *S. pennellii* introgressed region. At the chromosome 3 locus, 5 markers in total were used for genotyping: three were located in the region shared between IL3-3 and IL3-4, and one each was located in the region exclusive to either IL3-3 or IL3-4. Each individual was scored as a homozygous *S. lycopersicum* (LL), heterozygous (LP), or homozygous *S. pennellii* (PP) genotype for each chromosomal region based on these marker genotypes. During the generation of DILs, recombination events could occur within the introgression *S. pennellii* region within each line, so scoring LL, LP, and PP individuals at each locus required some additional criteria. In particular, individuals were classified into genotypic categories based on the identity (L vs. P) of the majority of markers scored within each target regions. For example on IL3-3 if three of the four markers were PP and one marker was heterozygous (LP), this individual was scored as double homozygous for *S. pennellii* (PP) at this chromosomal region. For IL 12 -3, if the three markers within the introgressed region disagreed, marker 74.00 was used as the tiebreaker as this marker is physically close to the genomic location of HT-A and HT-B. For each DIL population, genotype frequencies at both loci (e.g. chromosome 3 and 12) were determined by combining data from the two chromosomal regions to calculate observed two-locus genotype frequencies.

### Quantifying the presence and speed of UI

To determine pollen tube growth phenotypes in each of our NIL and DIL lines, and therefore the phenotypic expression of UI, we used an assay in which pollen is manually applied to a target stigma, allowed to germinate and grow in styles for 24 hours, and then styles are fixed and stained to visualize and measure the extent of pollen tube growth (Figure 1). Evidence of UI is demonstrated via rejection of pollen tubes in the female reproductive tract (style), quantified in terms of the proportional distance of pollen tube growth out of the total style length—a value that can vary between zero (very rapid rejection at the top of the style) to 1 (complete pollen tube growth down the entire length of the style).

**Figure 1.**
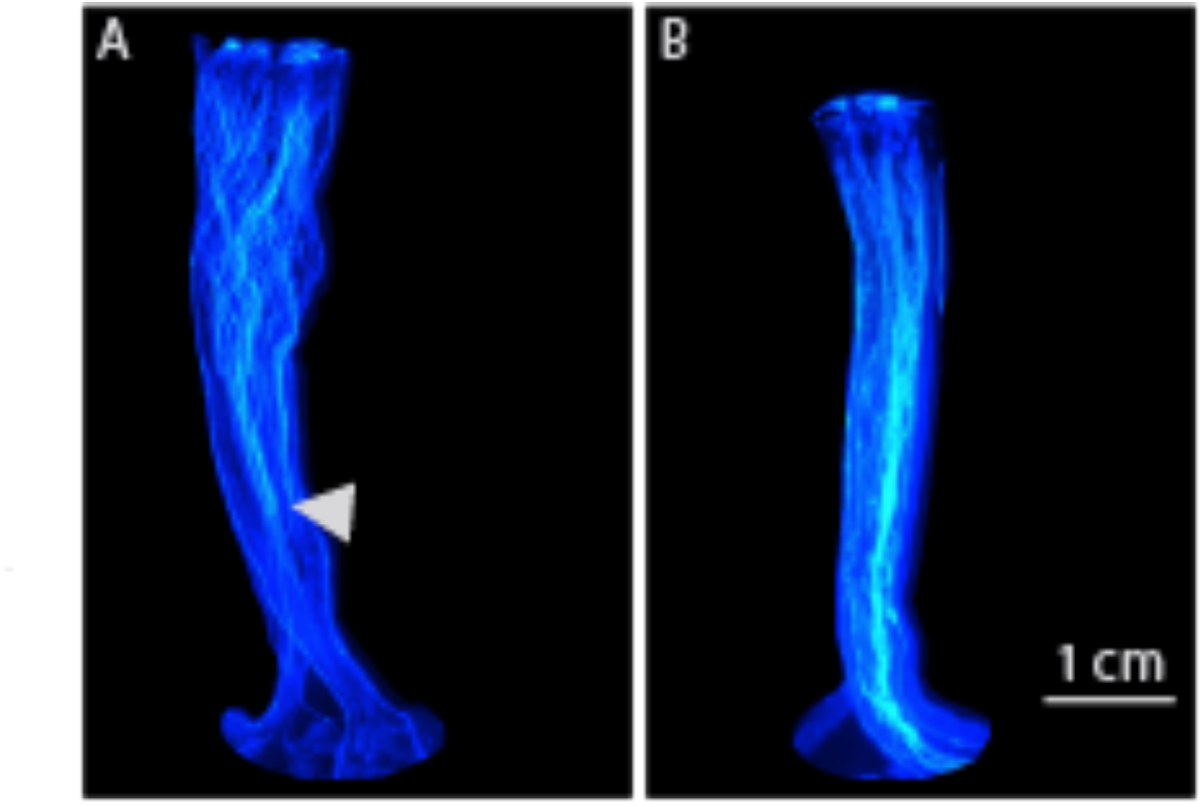
A) Representative images of the presence (A) and absence (B) of a UI pollen rejection response in the pistil. The white bar is the 1 cm scale used for measuring each style. A) Here, UI is illustrated with the phenotype observed in DIL 3-4 × 12-3, in which pollen rejection occurs approximately three-fourths down the length of the style. The arrowhead indicates where the majority of pollen tubes halt within the female reproductive tract (pistil) in this genotype. B) Compatible cross in which pollen tubes successfully reach the ovary, illustrated with the phenotype observed for NIL 3-4.

For each assay, an unopened bud was emasculated (1 day prior to opening) by removing the entire anther cone using hand forceps. Hand pollination was performed the following day, using *S. lycopersicum* (accession LA3475) pollen. This accession is the *S. lycopersicum* genomic background in our NILs and DILs; its pollen is expected to be rejected in a style that has an *S. pennellii*-derived genotype sufficient to mount a UI response. At 24 hours post-pollination, styles were collected into a 3:1 mixture of 95% EtOH:Glacial Acetic Acid in individual eppendorf tubes, and stored in the -20 freezer until imaging. 24 hours is more than sufficient for normal pollen tube growth down the complete length of the style, except when a UI response has been mounted. Pollination protocols were identical for both our NIL and DIL lines. We assayed 3 – 5 styles (technical replicates) per biological individual (Table 1).

To score pollen tube growth phenotypes, collected styles were placed in 5M NaOH and allowed to soften for 20 – 24 hours. Following softening, styles were washed and stained using 200 μL of Aniline blue fluorochrome for 3.5 hours in the dark, as described previously (Bedinger *et al.*, 2011; Jewell, 2016). Styles were then imaged using EVOS FL microscope with the DAPI setting. Stained pollen tubes fluoresce under these conditions, allowing us to differentiate style tissue from pollen tubes, and therefore determine the extent of pollen tube growth in each style. Because styles are generally too long to be captured in a single image, multiple images of the style were taken at 4x magnification and then stitched together using the program AutoStitch (Brown and Lowe, 2007). Images were visualized for measurement using ImageJ (Schneider *et al.*, 2012). Measurements taken on each style included 1) length of the style, 2) the “front” of the pollen (where the majority of the pollen stops in the style), and 3) the five longest pollen tubes. The average of the five pollen tubes was taken as our measurement of absolute distance traveled within this style, and UI was quantified as the proportional distance of pollen tube growth out of the total style length.

### Statistical analyses

To determine whether our observed genotypic frequencies significantly deviated from expected genotype frequencies in each ‘DIL population’, we calculated the expected proportion of each genotype and then performed a binomial test with a Bonferroni correction (Table 2). For each pairwise combination of loci, we used the upper and lower 95% confidence intervals around the regression coefficient to verify that the differences for genotypic classes were significant. We then performed planned independent contrasts for two different types of comparisons. Specifically, we asked 1) are DILs different from their parental NILs in terms of pollen tube growth and 2) are DILs different from each other? All analyses were run in RStudio version 0.99 (RStudio Team, 2015).

### Data availability

Introgression lines are available from the Tomato Genetics Resources Center (tgrc.ucdavis.edu). Marker information and phenotype data are supplied as supplementary material to the paper (Supplemental Table 1 and 2).

## RESULTS

### Departure from Mendelian Segregation Ratios

We observed significant TRD in the DIL populations generated from crosses between different NILs (Table 2A, B, C), suggesting that there are interactions among alleles at these loci that specifically affect the likelihood of transmission of different introgression combinations. For example, a strong deviation in our IL3-3 and IL12-3 population is due to overrepresentation of homozygotes for *S. pennellii* alleles at both chromosomal regions, as well as overrepresentation of individuals that are heterozygous (LP) on chromosome 3 and homozygous (PP) on chromosome 12. This observed pattern of TRD is consistent with selection against *S. lycopersicum* alleles on a heterozygous F1 pistil (that has one allele at each of these two *S. pennellii* regions) (Figure 2). While some genotypes within the other DIL combinations deviate somewhat from the expected ratios (Table 2B and 2C), only two additional comparisons survived Bonferroni correction: in the DIL 12-3 × 3-4 combination, we found evidence of underrepresentation of homozygotes for *S. pennellii* (PP) alleles at chromosome 12, specifically when heterozygous (LP) on chromosome 3. Similarly, in DIL 1-1 × 12-3, genotypes homozygous for *S. pennellii* (PP) at chromosome 1 and heterozygous (LP) at chromosome 12 were significantly underrepresented.

**Figure 2.**
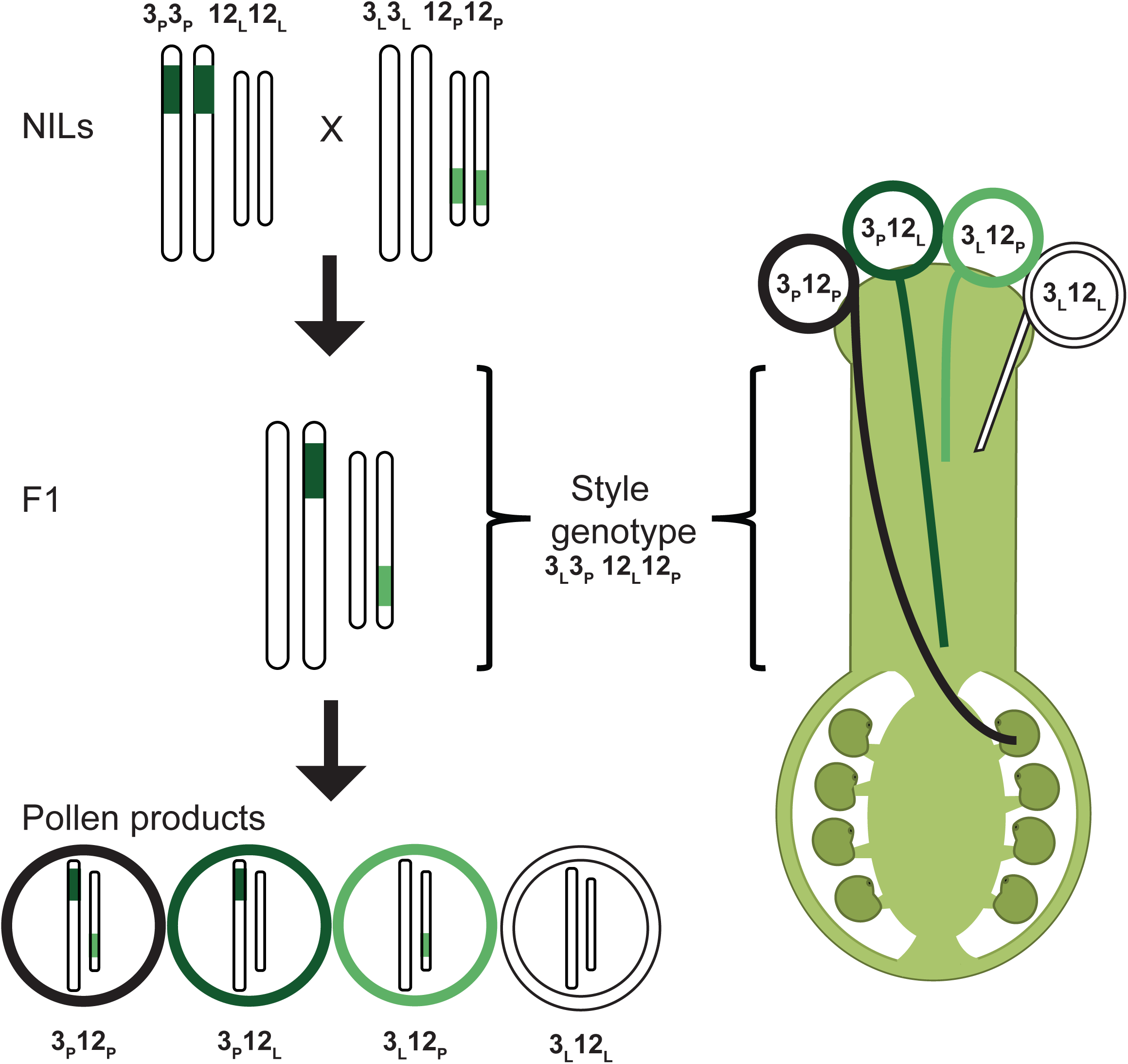
One model of gametophytic selection on different haploid pollen genotypes that could result in transmission ratio distortion among F2s. Two loci (*ui3.1* and *ui12.1*) are designated by whether they have alleles from *S. pennellii* (i.e., 3_P_ and 12_P_) or *S. lycopersicum* (i.e., 3_L_ and 12_L_). Left side: When two NILs are crossed (3_P_3_P_ 12_L_12_L_ × 3_L_3_L_12_P_12_P_), the F_1_ is heterozygous at the two regions (3_P_ 12_L_ 3_L_12_P_). Selfing this F_1_ produces four different haploid pollen genotypes: 3_L_12_L_, 3_L_12_p_, 3_p_12_L_, and 3_P_12_L_. Right side: Based on the genotype of our heterozygous F_1_ individual (3_P_ 12_L_ 3_L_12_P_) preferential pollen use, and stylar selection against specific pollen genotypes (e.g., 3_L_12_L_, 3_L_12_P_, and 3_P_12_L_), could generate deviations from expected Mendelian ratios.

**Table 2A.** Observed and expected (in parentheses) genotype frequencies for the crosses between chromosome region 12-3 (female) and chromosome region 3-3 (male) deviate from expected values, *X^2^* (8, N = 26) = 51.217, p-value = 2.38e-08. Genotypes that are overrepresented are indicated by +, after Bonferroni correction.

| Genotype at IL 12-3 |  |  |  |
| --- | --- | --- | --- |
| Genotype at IL 3-3 | LL | LP | PP |
| LL | 1 (1.63) | 1 (3.25) | 1 (1.63) |
| LP | 0 (3.25) | 2 (6.5) | 13 (3.25)<sup>+</sup> |
| PP | 1 (1.63) | 1 (3.25) | 6 (1.63)<sup>+</sup> |

**Table 2B.** Observed and expected (in parentheses) genotype frequencies for the crosses between chromosome region 3-4 (female) and chromosome region 12-3 (male) deviate from expected values *X^2^* (8, N= 117) = 25.86, p-value = 0.001103. Genotypes that are underrepresented are indicated by −, after Bonferroni correction.

| Genotype at IL 3-4 |  |  |  |
| --- | --- | --- | --- |
| Genotype at IL 12-3 | LL | LP | PP |
| LL | 14 (7.31) | 15 (14.63) | 11 (7.31) |
| LP | 18 (14.63) | 23 (29.25) | 23 (14.63) |
| PP | 4 (7.31) | 3 (14.63)<sup>-</sup> | 6 (7.31) |

**Table 2C.** Observed and expected (in parentheses) genotype frequencies for the crosses between chromosome region 12-3 (female) and chromosome region 1-1 (male) deviate from expected values, *X^2^* (8, N= 67) = 19.649, p-value = 0.0012. Genotypes that are underrepresented are indicated by −, after Bonferroni correction.

| Genotype at IL 12-3 |  |  |  |
| --- | --- | --- | --- |
| Genotype at IL 1-1 | LL | LP | PP |
| LL | 9 (4.18) | 13 (8.375) | 7 (4.18) |
| LP | 10 (8.375) | 17 (16.75) | 5 (8.375) |
| PP | 2 (4.18) | 1 (8.375)<sup>-</sup> | 3 (4.18) |

### UI phenotypes are observed in pairwise genotype combinations of ui3.1 with ui12.1

We found that, while individual NIL lines showed no UI response, several DIL genotype combinations exhibited significant UI (Supplemental Figure 2). The proportion of pollen tube growth down the style in the NILs ranged from 0.93 – 0.98, with confidence intervals that overlapped 1.00, indicating pollen tubes that have grown the entire length of the style and reached the ovary (Table 1, Figure 3). In addition, the 95% confidence intervals on each NIL mean overlap, consistent with no differences among NILs in their UI phenotype (Figure 3). In comparison, mean pollen tube growth was generally reduced in the DIL genotypes and more highly variable between them (0.54 – 0.85; Table 1). Of these DIL combinations, two show evidence for significant reductions in proportional pollen tube growth, consistent with a quantitative UI response; both these DILs involve combinations of *S. pennellii* alleles on chromosomes 3 and 12 (i.e. IL3-3 or IL3-4 with IL 12-3). Planned independent contrasts confirmed that DILs which combine *S. pennellii* alleles on chromosomes 3 and 12 were significantly different from their respective NIL parental genotypes. (DIL12-3 × 3-3: F_1, 6_ = 15.21; P = 0.0007, DIL 3-4 × 12-3: F_1, 6_ = 30.21; P = 1.6e-05). In comparison, the DIL combining *S. pennellii* alleles on chromosomes 1 and 12 was not significantly different than its parental NILs (F_1, 6_ = 2.226; P = 0.1500), and its 95% confidence intervals overlapped with the parental NIL genotypes, consistent with no significant interaction giving rise to a UI response in this DIL.

**Figure 3.**
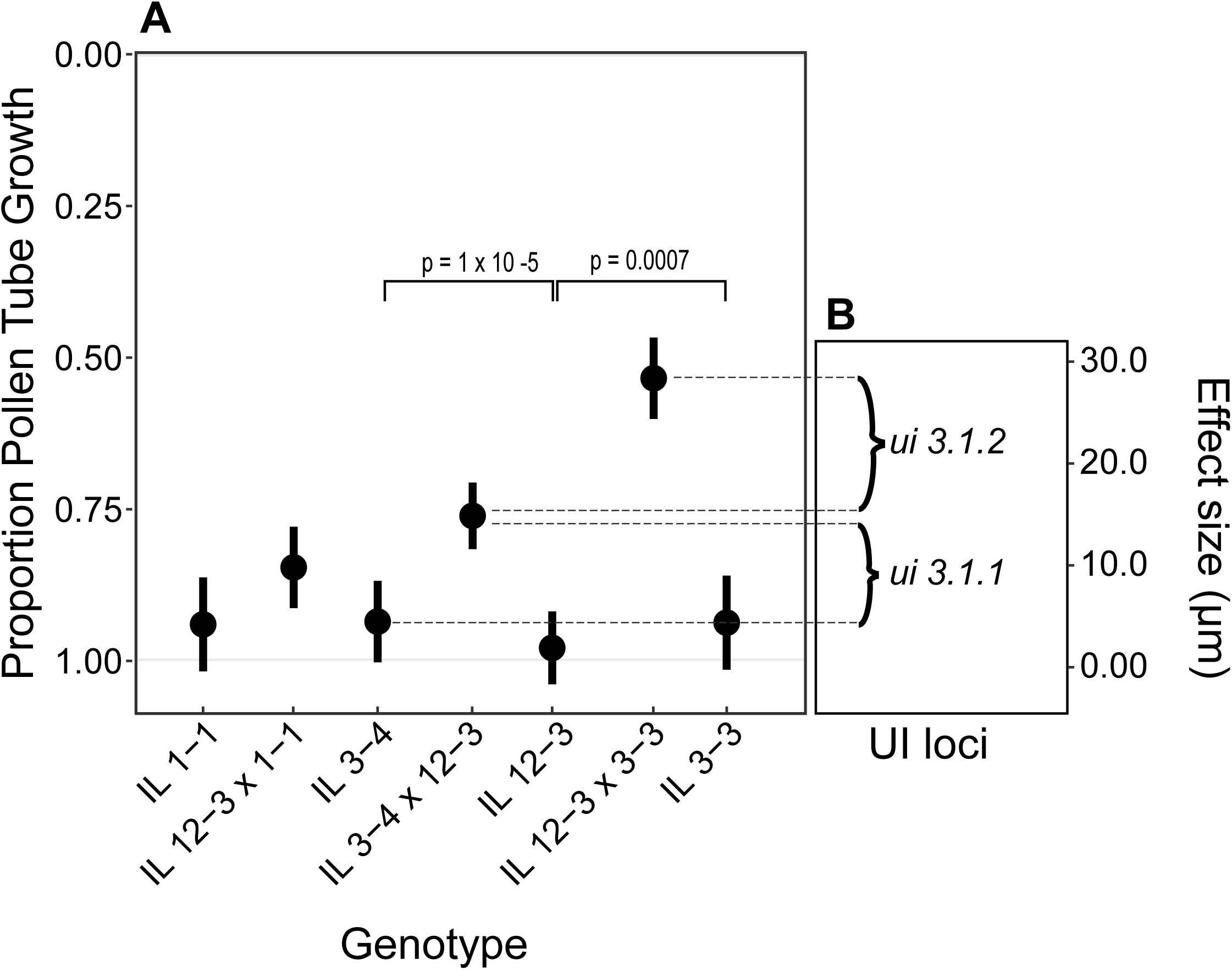
Proportion of pollen tube growth by genotypic class. A) Mean value with upper and lower 95% confidence intervals for distance traveled through the length of the style for each genotypic class. The top of the graph (0) represents the stigma, where pollen is placed. The bottom of the graph (1.0) represents the ovary, which is at the base of the style. B) The estimated effect size of the two loci (*ui3.1.1, ui3.1.2*) inferred to underlie *ui3.1* (see text).

### Patterns of pollen rejection suggest two loci underlie UI QTL on chromosome 3

In addition to displaying significant UI phenotypes, we also found that the quantitative strength of UI differed significantly between DILs with chromosome 3 *S. pennellii* alleles, depending upon which specific chromosome 3 introgression they contained (12-3 × 3-3 versus 34 × 12-3: F_1, 6_ = 12.835; P = 0.0017). In particular, DIL individuals carrying the *S. pennellii* alleles from IL3-3 showed a more rapid UI response (pollen tube growth arrests half way down the style) in comparison to DIL individuals carrying the *S. pennellii* alleles from IL3-4 (Figure 3A). These alternative DILs carry chromosome 3 introgressions that overlap by 44 cM (Supplemental Figure 1) as well as unique *S. pennellii* regions of 12 cM and 14 cM in IL3-3 and IL3-4 respectively, so the differences in UI phenotype between them could be explained by allelic differences within these unique regions. While there are several possible genetic interpretations of our observed patterns, the most parsimonious explanation requires only two loci at which *S. pennellii* alleles contribute to the quantitative expression of UI (in combination with *ui12.1*). The first locus (*ui3.1.1*) is contained within the chromosomal region shared between IL3-3 and IL3-4 and contributes an average effect of 0.19 micrometers to the quantitative strength of UI (see Figure 3B). In addition, the IL3-3 region contains a second locus, which, in combination with *ui3.1.1* and *ui12.1*, contributes an additional average effect of 0.23 micrometers to the quantitative strength of UI (see Figure 3B) when homozygous for *S. pennellii* alleles.

## DISCUSSION

While data suggest that there might be substantial overlap between molecular mechanisms of SI and UI, including both *S-RNase*-dependent or -independent mechanisms, several important aspects of the genetics of pistil-side UI remain unclear, including the minimum number of factors sufficient to express *S-RNase*-independent UI, the degree of overlap with molecular mechanisms underpinning *S-RNase*-dependent mechanisms, and therefore the level of redundancy between alternative mechanisms underlying these important postmating mechanisms of female mate choice. Here, our goals were to evaluate the minimum number of loci required to express pistil-side interspecific *S-RNase*-independent UI and evaluate the relative contribution of three different chromosomal regions to this phenotype. To do so, we examined the individual and combined (epistatic) effects of three candidate loci, pyramided as double introgression lines, on the expression of pistil-side pollen rejection. We find evidence that factors on both chromosomes 3 and 12 are jointly required for the expression of *S-RNase*-independent UI, whereas these loci have no effect individually. In addition, we find evidence consistent with gametophytic selection against certain genotypes, in the form of transmission ratio distortion. Together, these results suggest a strong role for the joint (epistatic) action of relatively few loci in determining the expression of pollen-pistil compatibility in this system.

### Two loci are jointly required to express S-RNase independent UI between species

Our findings clearly support a strong role for interactions between >1 molecular factor in the expression of *S-RNase*-independent UI. Specifically, we show that *S. pennellii* alleles from both chromosome 3 and 12 are necessary (and sufficient) for the expression of quantitative pistil-side UI against SC *S. lycopersicum* pollen. This confirms the general expectation that pollen recognition and rejection requires coordinated molecular interactions between several proteins, consistent with other kinds of molecular recognition and rejection mechanisms (McClure *et al.*, 1999; McClure *et al.*, 2000), but also indicates that the products of relatively few loci are sufficient to mount an *S-RNase*-independent UI response. Our study was designed based on previous mapping studies of UI in *Solanum* which identified main effect loci for UI on chromosomes 3, 12, and/or 1 (Bernacchi and Tanksley, 1997). Other analyses have confirmed that *ui1.1* is associated with the presence/absence of functional pistil-side *S-RNase*, at least between *Solanum* lineages in which one genotype is SI. The involvement of *S-RNase* for interspecific pollen rejection was confirmed via transformation of SC *Nicotiana* species in which pollen rejection occurred from non transformed SC *Nicotiana* species demonstrating that *S-RNase* is sufficient for UI ((Murfett *et al.*, 1996). Nonetheless, because the *S*-locus is a large and genetically complex chromosomal region, additional loci within this region might also contribute to *S-RNase*-independent UI among genotypes that lack *S-RNase* but still show UI phenotypes. Our results, however, do not support the involvement of additional pistil-side loci at *ui1.1* in UI rejection in this particular species pair; we detected no additional effect of *S. pennellii* alleles at *ui1.1* between our two genotypes, both of which lack *S-RNase* function.

In contrast to *ui1.1*, our results confirm that *S-RNase*-independent UI is the joint product of *S. pennellii* alleles at chromosomes 3 and 12. Molecular analyses in other *Solanum* species pairs indicate that pistil-side HT protein contributes to the effect associated with *ui12.1* between lineages showing UI. Requirement of HT protein was demonstrated for pollen rejection from *N. plumbaginifolia* (Hancock *et al.*, 2003) and within *Solanum* one of the tandemly duplicated HT proteins (HT-A) was detected and expressed for a number of species implicating its function in UI (Covey *et al.*, 2010). In addition, a QTL analysis between *S. lycopersicum* and a different (SI) genotype of *S. pennellii* (Jewell, 2016) indicated that presence/absence of HT protein was significantly associated with the strength of the UI response in a segregating F2 population. Although we do not have equivalently direct data on the molecular underpinnings of *ui12.1* here, based on these other studies our working hypothesis is that the effect of IL12-3 on UI is partly or solely due to *S. pennellii* alleles at HT.

In comparison to *ui1.1* and *ui12.1*, the molecular loci underpinning *ui3.1* remain unknown, although Jewell (2016) identified several potential genes, and three especially strong candidates, for this locus by combining additional genomic and gene expression data (Pease *et al.*, 2016) with the mapped location of *ui3.1* in that study. Interestingly, our findings here suggest that *ui3.1* is more complex than revealed in that and other mapping experiments. QTL analysis has known limitations in terms of identifying number, location, and individual effects of loci because detection depends on the heritability of the trait, the size of the segregating population, and the density of genetic markers (cite). Accordingly, the observation that single QTL can resolve into more than one underlying locus is not uncommon (Mackay *et al.*, 2009), especially when the confidence intervals on this locus are broad. In this case, *ui3.1* was mapped to a region of ~100 cM between *S. pennellii* and *S. lycopersicum* (Jewell, 2016) and ~10 cM between *S. habrochaites* and *S. lycopersicum*, (Bernacchi and Tanksley, 1997), both of which could easily harbor >1 contributing locus.

Given this, currently the most parsimonious inference from our observations is that (at least) two loci contribute additively to quantitative UI expression at *ui3.1*, one located in the genomic region overlapping between IL3-3 and IL3-4, and one in the region unique to IL3-3 (see Results, and Figure 3B). Nonetheless, there are other alternative interpretations of our observations. For example, tomato chromosomes are known to be enriched for pericentromeric heterochromatin (Wang *et al.*, 2006; Tanksley *et al.*, 1992). Marker delineated breakpoints indicate that IL3-3 overlaps the centromeric region, and therefore contains centromeric heterochromatin from *S. pennellii* that matches the species origin of the rest of the introgression. In comparison, IL3-4 does not contain the *S. pennellii* centromeric region. Given this, it’s possible that the phenotypic difference between IL3-3 and IL3-4 is instead due to position effects at *ui3.1* that are associated with proximity to conspecific (IL3-3) versus heterospecific (IL3-4) centromeric heterochromatin. Interestingly, this still implies that two loci are involved in the phenotypic patterns we observed at *ui3.1*, just that the second ‘locus’ is the regulatory environment associated with species differences in the pericentromeric region.

Regardless, in terms of our goals to evaluate the minimum number of loci required to express pistil-side *S-RNase*-independent UI, and evaluate the relative contribution of three different chromosomal regions to this phenotype, our data indicate that two to three loci at two genomic locations—on chromosomes 3 and 12—are jointly required and sufficient to express this important postmating interspecific barrier.

### Source of Transmission Ratio Distortion

We detected evidence of transmission ratio distortion during the generation of our double introgression lines. One interpretation is that these distorted genotypes are due to gametophytic selection against particular haploid pollen genotypes in the F1 (doubly heterozygous) style in each case. Genetic differences among male gametophytes could result in either differential gametophytic selection or competition, both of which could potentially influence genotype frequencies in the next generation (Snow and Mazer, 1988). For example, gametophytic selection could change the probability of fertilization based on genetic differences expressed in the male gametophyte according to a simple model illustrated in Figure 2. In this case, pollen that carries both *S. pennellii* alleles for IL 3-3 and IL 12-3 has a growth or persistence advantage in the heterozygous F1 pistil whereas pollen carrying *S. lycopersicum* alleles is preferentially selected against (Figure 2). This model implies that S. pennellii-derived proteins in the style are able to differentially recognize and reject pollen that lacks *S. pennellii* alleles at *ui3.1* and *ui12.1.* This is intriguing because these loci are expected to have pistil-side functions but are not necessarily expected to mediate pollen-side involvement in UI. In contrast, known loci that influence pollen-side expression of UI are located on chromosomes 1, 6, and 10 (Li and Chetelat, 2015; Li and Chetelat, 2010; Li *et al.*, 2010; Chetelat and DeVerna, 1991) although these observations do not exclude the potential involvement of additional loci on other chromosomes. We note also that dissimilar patterns of TRD are observed in DIL populations from two other NIL combinations, where some genotypes with *S. pennellii* alleles are significantly underrepresented (Results). In these later cases, however, we cannot exclude the possibility that pollen carrying heterospecific chromosomal segments is simply less viable or less competitive than ‘pure’ S. *lycopersicum* pollen (i.e. that TRD is due to reduced hybrid male fertility in these genotypes). In addition, patterns of TRD can also be due to other complex causes such as dysfunction (hybrid inviability) in both male and female gametes that results in TRD due to differential (sex independent) gamete survival (Koide *et al.*, 2008) or to differential survival of post-fertilization hybrid zygotes. Therefore, although gametophytic selection might explain part or all of the patterns of TRD we observed, other complex causes are also possible.

### Unilateral Incompatibly and Reproductive Isolation

The active recognition and rejection of heterospecific pollen tubes growing within the pistil is a form of postmating cryptic female choice against heterospecific mates. In the Solanaceae, UI is a particularly common reproductive isolating barrier, especially in genera that have both SI and SC species (Bedinger *et al.*, 2011; Lewis and Crowe, 1958). Dissecting the mechanisms that contribute to UI can therefore provide insight into the expression of this reproductive barrier among lineages. Here we have shown that relatively few loci are sufficient to express UI. Moreover, these pistil-side mechanisms likely share genetic components with self-incompatibility mechanisms within species. Previous work has shown that *S-RNase* necessary for SI is also a key pistil-side contributor to UI (Murfett *et al.*, 1996), as is HT (McClure *et al.*, 1999). Moreover, HT appears to contribute to both *S-RNase* dependent and *S-RNase* independent mechanisms of UI among some species (Covey *et al.*, 2010; Tovar-Méndez *et al.*, 2016). Given these dual functions, it is plausible that the molecular machinery for mounting a UI response is already present in the styles of SI species, and its phenotypic expression merely requires encountering heterospecific SC pollen. Nonetheless, the existence of several redundant mechanisms (i.e. *S-RNase* independent and dependent mechanisms) of UI suggests that this heterospecific barrier is not merely a pleiotropic by-product of stylar competence for SI. Indeed, it is possible that the molecular mechanism(s) underlying *ui3.1* are not shared with SI recognition and rejection, consistent with independently selected and maintained mechanisms specifically for UI. Evaluating this possibility awaits further fine-mapping and functional characterization of this locus (or loci) in the future.

## ACKNOWLEDGMENTS

The authors thank Ashley Huh and C.J. Jewell for data collection and assistance with experimental material. The authors would also like to thank Dean Castillo for advice on statistical analyses. We also thank the Indiana University Bloomington greenhouse staff. Finally, the authors would like to thank X # of reviewers for comments that greatly improved the manuscript. This research was funded by NSF grant MCB-1127059.

## Supplementary Material

**Supplemental Table 1.** Near isogenic line accessions used in this study. Accession numbers are from the Tomato Genetics Resource Center (tgrc.ucdavis.edu).

| Accession | Introgression Line | n* |
| --- | --- | --- |
| LA 4028 | IL 1-1 | 3 |
| LA 3488 | IL 3-3 | 3 |
| LA 4046 | IL 3-4 | 4 |
| LA 4100 | IL 12-3 | 5 |

\* Number of biological replicates per accession

**Supplemental Table 2.** CAPs markers used for genotyping with marker ID based on Solgenomics markers.

| Ch | cM | Marker ID | Enzyme* | PCR Fragment | L Fragment | P Fragment |
| --- | --- | --- | --- | --- | --- | --- |
| 1 | 30.20 | C2_At3g06580 | *DpnII* | 300 | 170 | 270 |
| 1 | 40.00 | C2_At4g30890 | *TaqI* | 600 | 300+200 | 350+200 |
| 1 | 58.00 | C2_At1g48050 | *TaqI* | 920 | 920 | 750 |
| 3 | 72.40 | C2_At3g02910 | *TaqI* | 450 | 150 | 325 |
| 3 | 101.50 | C2_At3g47990 | *HaeIII* | 1100 | 1100 | 780+320 |
| 3 | 113.00 | C2_At1g74520 | *RsaI* | 1350 | 410+390 | 1350 |
| 3 | 120.50 | C2_At5g62440 | *DraI* | 950 | 700 | 570 |
| 3 | 128.50 | C2_At3g47640 | *XbaI* | 850 | 850 | 550+300 |
| 12 | 52.00 | T1736 | LP* | - | 1200 | 1050 |
| 12 | 74.00 | T0801 | *AccI* | 589 | 400 | 600 |

Ch = chromosome, cM = centiMorgans, \* LP is a linked polymorphism (i.e. size difference noticeable after first PCR reaction so no digestion via restriction enzymes necessary), L Frag = Expected size if double homozygous *S. lycopersicum*, P Frag = Expected Size if *S. pennellii*

Supplemental Figure Legend

Supplemental Figure 1. Representative images of the softened and stained pollinated styles, showing proportion of pollen tube growth towards the ovary for each genotype evaluated. White scale bars are 1000 μm.

Supplemental Figure 2. The relative locations, and overlapping and non-overlapping introgressed regions, for IL 3-3 and IL 3-4 on chromosome 3. Markers locations used for genotyping are illustrated on the right; a single marker identifies either IL 3-3 (marker 72.40) or IL 3-4 (marker 128.50), while the three intervening markers fall within the overlapping region. Map generated based on map distances obtained from Solgenomics.net.

